# Defense systems are pervasive across chromosomally integrated mobile genetic elements and are inversely correlated to virulence and antimicrobial resistance

**DOI:** 10.1101/2022.11.18.517082

**Authors:** João Botelho

**Affiliations:** Centro de Biotecnología y Genómica de Plantas (CBGP), Universidad Politécnica de Madrid (UPM) - Instituto Nacional de Investigación y Tecnología Agraria y Alimentaria (INIA-CSIC), Madrid, Spain

## Abstract

Mobile genetic elements (MGEs) are key promoters of microbial evolution. These elements can be located extrachromosomally or integrated into the chromosome. Well-known examples of chromosomally integrated MGEs (ciMGEs) are integrative and conjugative/mobilizable elements (ICEs and IMEs), and most studies to date have focused on the biological mechanisms that shape their lifestyle. It is crucial to profile the diversity and understand their distribution across the microbial community, as the number of genome sequences increases exponentially. Herein, I scanned a collection of more than 20000 bacterial and archaeal non-redundant genomes and found over 13000 ciMGEs across multiple phyla, representing a massive increase in the number of ciMGEs available in public databases (<1000). Although ICEs are the most important ciMGEs for the accretion of defense systems, virulence, and antimicrobial resistance (AMR) genes, IMEs outnumbered ICEs. Moreover, defense systems, AMR, and virulence genes were negatively correlated in both ICEs and IMEs. Multiple ciMGEs form heterogeneous communities and challenge inter-phylum barriers. Finally, I observed that the functional landscape of ICEs was populated by uncharacterized proteins. Altogether, this study provides a comprehensive catalog of nucleotide sequences and associated metadata for ciMGEs from 34 phyla across the bacterial and archaeal domains.

## Introduction

Bacterial and archaeal genomes have evolved diverse defense systems to target endoparasites, such as bacteriophages, plasmids, and other types of mobile genetic elements (MGEs) that can be in conflict with their host cells (1). In fact, more than one class of defense systems are usually found in the same cell, nonrandomly clustered in so-called defense islands (2). Different defense systems can work together or independently to defend the host against foreign DNA (3, 4). Besides, a recent study found that antimicrobial resistance (AMR) genes and defense systems can cluster together in MGEs (5). Worryingly, this study found that phage infection itself spurs the excision of integrative and conjugative elements (ICEs) and subsequent transfer by conjugation, leading to the spread of AMR genes. Due to a high fitness cost and to the evolutionary arms race between defense systems and MGEs, these systems are expected to be prone to loss (6, 7), and in the long-term their co-occurrence with genes involved in mobilization is crucial for persistence. For example, two of the most well-known defense systems (toxin-antitoxin and restriction-modification) exploit MGEs for their spread, showing a distinct evolutionary history from their prokaryotic hosts (8). From an evolutionary viewpoint, it is tempting to propose that the presence of multiple defense systems may be more involved in the maintenance of MGEs than of the cell (1, 9). Besides, MGEs can use additional strategies to offset the short-term deleterious effects of their cargo genes, such as increased rates of conjugation (10, 11). Horizontal gene transfer (HGT) and gene loss have crucial roles in the patchy distribution of defense mechanisms and AMR genes at the species and strain level (12).

Pathogenicity islands are a group of chromosomally integrated MGEs (ciMGEs), including different types of bona fide MGEs such as ICEs, integrative and mobilizable elements (IMEs), and actinomycete ICEs (AICEs), among others. ICEs are transferred as linear single-stranded DNA and use a type-IV secretion system (T4SS) for conjugation (similar to the mechanism used by conjugative plasmids) (13). In Actinobacteria, however, some ICEs are delivered as double-stranded DNA following a distinct process (14). As so, these ICEs have been categorized as AICEs. While AICEs and ICEs encode an intact conjugation apparatus, and are thus self-transmissible, IMEs encode their own excision and integration module but lack a fully functional conjugative apparatus for autonomous transfer. In addition to ICEs, AICEs, and IMEs, there are *cis*-mobilizable elements (CIMEs), which are flanked by *attL* and *attR* recombination sites but lack conjugation or recombination genes (15). Still, IMEs and CIMEs can pirate conjugative plasmids and ICEs to promote their own dissemination (16). Both elements are a driving force for bacterial adaptation and evolution, at least in part due to the different cargo genes that may provide the host a selective advantage in specific environments (17).

Since phage predation selects for multiple defense systems that frequently cluster on mobilizable defense islands, I explore here if different ciMGE types (i.e., ICEs, IMEs, AICEs, and CIMEs) are important hotspots for the accretion of defense systems, helping to protect the host from superinfection by other MGEs. Considering the known evolutionary relationship MGEs have with defense systems (2, 4, 18–20), AMR (5, 20–22) and virulence genes (23), I also explore if these three functions are positively or negatively correlated across ciMGEs from multiple phyla. I found that i) IMEs and ICEs are widespread across different phyla; ii) ICEs are the most common ciMGEs for the accumulation of defense systems, AMR, and virulence genes; iii) these three functions are negatively correlated across ICEs and IMEs; and iv) ICEs and IMEs from multiple phyla share high genetic similarity and challenge interphylum barriers. Finally, this work provides a comprehensive resource compiling 13274 ciMGEs from Bacteria and Archaea with associated metadata extracted from 34 distinct phyla.

## Material and methods

### Genome collection

Bacterial genomes sequenced at the complete level (i.e., all replicons included inside the genomes, such as the chromosomes and extra-chromosomal elements, are fully assembled with gaps not exceeding ten ambiguous bases) were downloaded from NCBI RefSeq (08-07-2021) using ncbi-genome-download v0.3.0 (https://github.com/kblin/ncbi-genome-download). Archaeal genomes were downloaded using the same approach (28-06-2022). To examine the presence of redundant genomes, I used Assembly Dereplicator v0.1.0 (https://github.com/rrwick/Assembly-Dereplicator) with a Mash distance clustering threshold of 0.001 and a batch size of 25000. The classify workflow from GTDB-Tk v2.0.0 (24) was used to correct the taxonomy classification of the downloaded bacterial and archaeal genomes. Two genomes (RefSeq assembly accession numbers GCF_900660555.1 and GCF_002158865.1) were excluded, as these were flagged by the align module in GTDB-Tk as having an insufficient number of amino acids in the MSA.

### Identification of ciMGEs

I then used a custom python script to split all genome files into 45482 individual replicon files (i.e, chromosomal and extra-chromosomal replicons that are part of the genomes). A total of 22269 replicons with the word ‘plasmid’ in the fasta-headers were removed. To validate the separation of chromosomal and plasmid replicons, I compared their sequence length distributions using a density plot. The remaining 23213 chromosomal replicon files were used as input in ICEfinder (25) to look for ciMGEs (i.e., ICEs, IMEs, AICEs, and CIMEs). The ciMGEs carrying an integrase gene, a relaxase gene, and T4SS gene clusters are considered ICEs, while the elements without T4SS but with integrase, replication and the AICE translocation-related proteins are classified as AICEs. IMEs code for their own excision and integration. However, they lack an intact conjugative machinery for autonomous conjugative transfer. The ciMGEs were then extracted from the chromosomes and reannotated with Prokka v1.14.6 (26). The resulting protein files from Prokka annotation were used as input to search for relaxases against the MOBfamDB protein profiles (27) using hmmscan from hmmer v3.3.2 (28). To explore potential artifacts resulting from the integration of plasmids into the chromosome, the extracted ciMGEs were scanned against the PlasmidFinder database (473 sequences, 25-02-2023) (29) through abricate v1.0.1 (https://github.com/tseemann/abricate).

### Functional annotation and network analysis

The resulting proteins and gff files from Prokka annotation were used as input to search for defense systems with PADLOC v1.1.0 (DB v1.4.0) (30). AMRFinder v3.10.30 (31) was used to search for the presence of AMR genes across the ciMGEs. To look for virulence genes, I used the virulence factor database VFDB (32) (4327 genes, 27-06-2022) with abricate v1.0.1. The ciMGE dataset was then dereplicated with MMseqs2 v13.45111 (33) using 90% sequence identity and 80% coverage. To explore if similar ciMGEs were present in different taxa, I then used the ciMGEs that were clustered above the threshold with MMseqs2. Cluster of orthologous groups (COG) categories were searched with eggNOG-mapper v2.1.9 (DB v5.0.2) (34). Since no CIMEs were identified in this study, from here on ciMGEs types only refer to IMEs, ICEs, and AICEs. To estimate the pairwise distances between all ciMGE types, I reduced the dereplicated ciMGEs into sketches and compared the Jaccard index (JI) between pairs of ciMGEs using BinDash v 0.2.1 (35). Only results with a p-value of at most 0.05 were reported. By default, BinDash reduces each sample into 2048 hash values. Each ciMGE nucleotide sequence was converted to a set of 21-bp k-mers. The JI values were used as weights to plot the undirected and weighted network and the communities detection was performed with Infomap (36) and OSLOM v2.5 (37). Infomap was called 5 times in OSLOM, followed by the software’s cleanup procedures to the communities found by Infomap. The p-value (-t parameter) for community detection was set to 0.05, and the -singlet parameter was used to filter noisy nodes and to avoid assigning singletons to an existing community. I then used Cytoscape v3.9.1 (https://cytoscape.org/) to group the nodes according to the OSLOM communities identified in the first hierarchical level.

### Statistics

Comparisons between i) the number of defense systems, virulence, and AMR genes normalized to ciMGE size per phylum; ii) GC content deviation (GC host genome – GC ciMGE) across the different ciMGE types; iii) the number of cargo genes per ciMGE per phylum; iv) the COG counts normalized to ciMGE size were performed using the Kruskal-Wallis test, and the p-values adjusted using the Holm–Bonferroni method. Comparisons between the normalized number of COG categories between ICEs and IMEs were performed using the Wilcoxon test, and the p-values adjusted using the Holm–Bonferroni method. Correlation between defense systems, AMR, and virulence genes was assessed by the Spearman method. This method was also used to compute the correlation coefficient between the GC content and sequence length of ciMGEs and their host genome. Values above 0.05 were considered as non-significant (ns). We used the following convention for symbols indicating statistical significance: * for p <= 0.05, ** for p <= 0.01, *** for p <= 0.001, and **** for p <= 0.0001.

## Results

### IMEs and ICEs are widespread across multiple phyla

A total of 22334 complete genomes from 57 phyla (21897 genomes in 48 bacterial phyla and 437 genomes in 9 archaeal phyla, **Supplementary Table S1**) were used as input to search for ciMGEs. More than 80% of the downloaded genomes belong to three bacterial phyla: Proteobacteria, Firmicutes, and Actinobacteriota (n=11843, 4706, and 2226, respectively, **Supplementary Figure S1**). In order to focus exclusively on MGEs integrated into the chromosome, plasmid replicons were first removed. While chromosomes follow a bell-shaped distribution, plasmid replicons are instead right-skewed (**Supplementary Figure S2**) (38). The search resulted in the identification of 13274 ciMGEs, including 6331 IMEs, 5869 ICEs, 1061 AICEs, and 13 elements annotated as putative conjugative regions. The latter corresponds to regions detected by ICEfinder in small replicons (≤ 1Mb, **Supplementary Table S2**) with an integrase and a qualified T4SS but no relaxase nor a type-IV coupling protein. These 13274 ciMGEs were extracted from a total of 34 phyla (13258 ciMGEs in 8000 bacterial genomes and 16 ciMGEs in 16 archaeal genomes, **Supplementary Table S2**). This means that 36.5% of bacterial genomes (8000/21897) carry at least one ciMGE, while only 3.7% of archaeal genomes (16/437) carry at least one ciMGE. As expected, the most prevalent phyla (Proteobacteria, Firmicutes, Actinobacteriota, Bacteroidota, Campylobacterota, and Firmicutes A) had a larger absolute number of IMEs and ICEs, with each phyla carrying more than 100 ICEs and IMEs (**Supplementary Tables S1 and S2**). IMEs were the most frequently identified ciMGE across our dataset (**Figures 1A and 1B)**, both in bacteria and archaea (n=6319 and 12, respectively, across a total of 4749 genomes), and the element type with the widest dispersion across different phyla (32 out of the 57, **Supplementary Table S2**). ICEs were also widely disseminated in bacterial and archaeal genomes (n=5865 and 4, respectively, across a total of 4328 genomes), and across 25 phyla. While these ciMGEs were found in both domains, AICEs were only found in bacteria (n=1061 in 519 genomes). In fact, AICEs were constrained to particular phyla (mostly in Actinobacteriota, and rarely in Firmicutes and Proteobacteria). When comparing with the number of ciMGEs deposited in ICEberg with nucleotide sequences available (n=718 ICEs, 111 IMEs, and 50 AICEs), this analysis considerably expands the repertoire of these elements.

**Figure 1.**
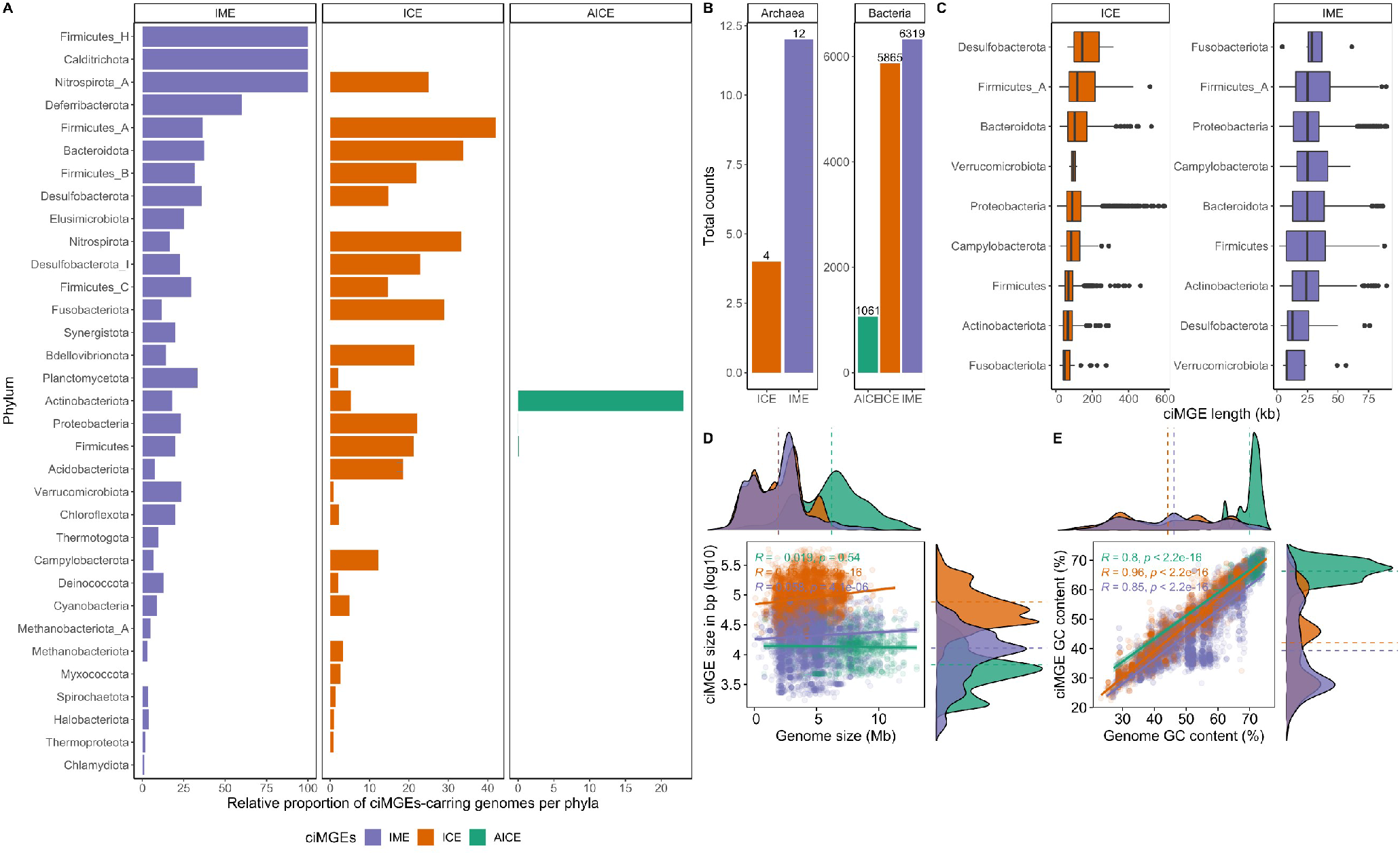
Distribution of ciMGEs across Bacteria and Archaea. **A)** Relative proportion of IME-, ICE-, and AICE-carrying genomes per phyla. The single proteobacterial genome with an AICE is not visible in the figure due to the low relative proportion (1 genome out of 11843 Proteobacteria). Phylum in the y-axis are listed in descending order. **B)** Total counts of ICEs, IMEs, and AICEs across bacterial and archaeal genomes. **C)** Boxplots showing the variation in ICE and IME’ size across multiple phyla. Phyla in the y-axis are listed in descending order of the median value for each boxplot. Only phyla with more than 30 ciMGEs are shown. **D)** Scatter plot with marginal density plots comparing the ciMGE size in bp and log10 scaled and the size of the host genome in Mb. Mean values for each ciMGE type are represented by dashed lines in each density plot. **E)** Scatter plot with marginal density plots comparing the ciMGE GC content (%) and the GC content of the host genome. Mean values for each ciMGE type are represented by dashed lines in each density plot. Bars, boxplots, density curves, regression lines, dashed lines, and correlation coefficients are coloured according to the ciMGE type.

I then explored the distribution of the sequence length and GC content from the ciMGEs identified in this study. When ICE and IME sizes were broken down by phyla, it was feasible to see that Desulfobacterota had the largest ICEs and Fusobacteriota had the largest IMEs (**Figure 1C**). Overall, IMEs follow a normal distribution for the sequence length, with a mean size of 27kb, while AICEs have asymmetrical distributions, with mean sizes of 17kb and 109kb, respectively (**Figure 1D**). Unsurprisingly, given the presence of a full conjugative type IV secretion system on ICEs, these elements tend to be larger than IMEs and AICEs. No correlation was found between the ICEs/IMEs and host genome size (R = 0.15 and p-value < 2.2e-16, R = 0.058 and p-value = 4.1e-06, respectively).When it comes to the GC content, while AICEs again have asymmetrical distributions and a mean GC content of 67%, IMEs follow a bimodal distribution and ICEs a multimodal distribution, with mean GC contents of 46% and 48%, respectively (**Figure 1E**). Genome GC content strongly correlates to ICEs, IMEs, and AICEs’ GC content, following a trend similar to plasmids (39) (R ≥0.8, p-value < 2.2e-16). No correlation was observed between ICE/IME size and GC content (R = 0.17 and p-value < 2.2e-16, R = 0.035 and p-value = 0.0052, respectively), nor between AICE size and GC content (R = −0.15 and p-value = 5.8e-07, **Supplementary Figure S3A**). When comparing the ciMGEs’ GC content with that of the host genome, IMEs’ GC deviation from that of their hosts was significantly higher than the difference observed for ICEs and AICEs (**Supplementary Figure S3B,** p-value < 2.2e-16). Altogether, these results show that IMEs outnumber ICEs, the GC content of both elements is strongly correlated to that of their host, and both elements are widespread across multiple phyla.

### Defense systems, AMR genes, and virulence genes are preferentially located in ICEs

Known defense systems are pervasive across bacterial ciMGEs (**Supplementary Table S2**). There are 4588 ciMGEs with at least one defense system gene: a total of 26876 hits in 4586 bacterial ciMGEs (34,6%), and only 32 hits in 2 archaeal ciMGEs (12,5%). These hits are dispersed across 23 phyla. Even though the total number of ICEs found in this study is smaller than that of IMEs (5869 vs 6331), these elements are the most important hotspots for the accretion of defense systems across ciMGEs. There are 2519 ICEs with a total of 15897 defense system genes, 1965 IMEs and 103 AICEs with a total of 10619 and 388 genes, respectively. Nearly half of ICEs carry at least one defense system (42,9%), while nearly a third of IMEs carry at least one of these systems (31,0%). Given that ICEs and IMEs have a diverse range of sizes across different phyla (**Figure 1D**), I corrected the number of defense system genes for the size of the carrying ciMGE and found that IMEs across major phyla carry more defense systems per kb than do ICEs (**Figure 2A**). Restriction-modification systems, abortive infection, and CBASS are among the defense systems most frequently found across ICEs, while restriction-modification systems, CBASS, and Thoeris represent a large share of the defense systems found on IMEs (**Figure 2B** and **Supplementary Table S3**). Multiple DNA modification systems (DMS) were found across these elements. The DMS model was developed by PADLOC to select potential defense systems where there are two or more genes from modification-based systems such as restriction-modification, BREX, and DISARM. Hits found by the DMS model will require manual curation. A total of 21 defense systems are found on AICEs from Actinobacteriota, including restriction-modification systems, ietAS, and BREX.

**Figure 2.**
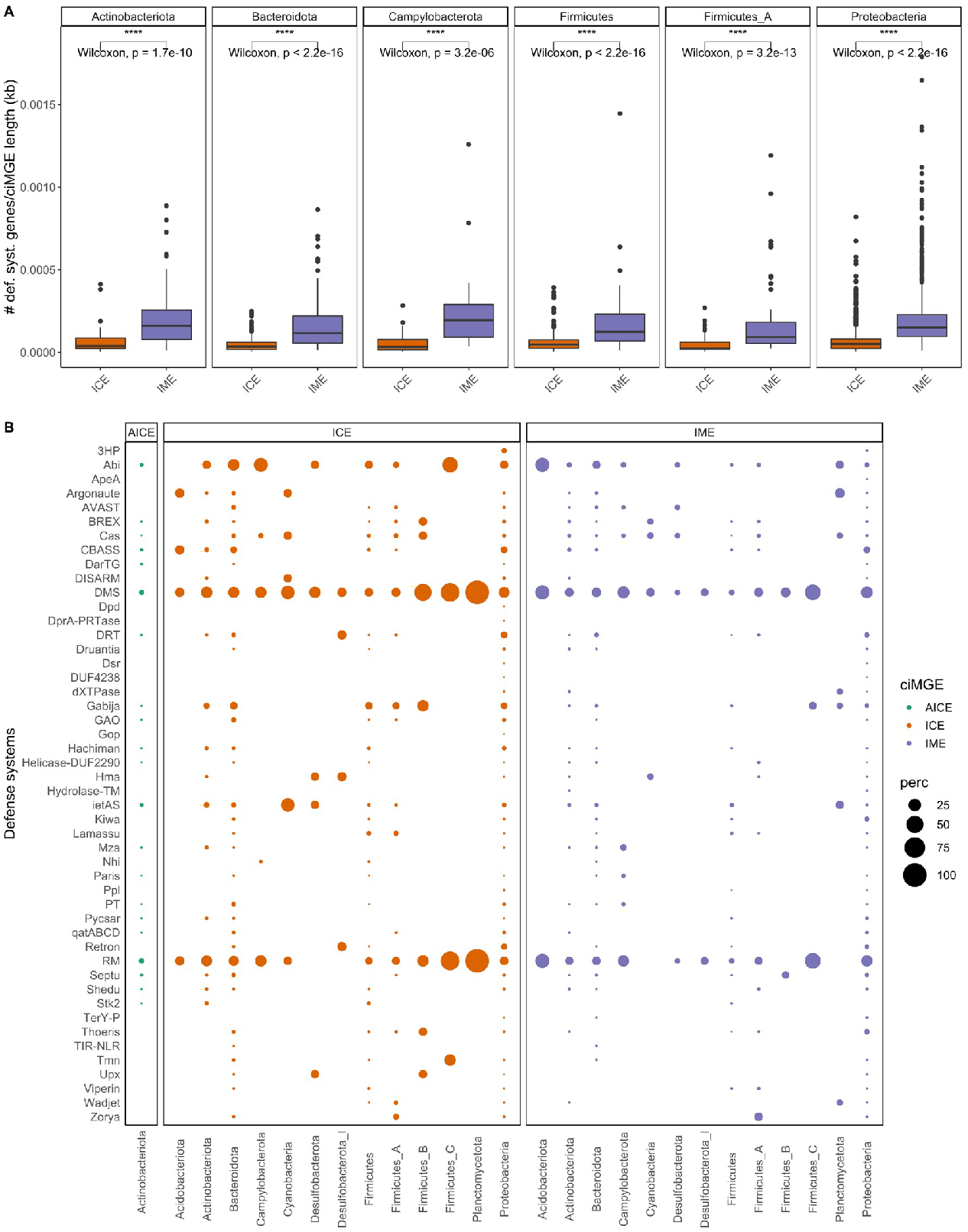
Defense systems are widespread across multiple phyla. **A)** Boxplots showing the variation between the number of defense systems normalized by ICEs and IMEs’ size (kb). Only phyla with more than 10 ciMGEs carrying defense systems are shown. Only statistically significant comparisons are shown above the boxplots. The following convention was used for symbols indicating statistical significance: * for p <= 0.05, ** for p <= 0.01, *** for p <= 0.001, and **** for p <= 0.0001. **B)** Distribution of defense system genes across multiple phyla. Phyla in the x-axis are listed from left to right in alphabetical order. Only phyla with more than 5 different defense systems are shown. The size of the circles is proportional to the percentage of particular defense systems per ciMGE type per phylum. Boxplots and circles are coloured according to the ciMGE type.

Known virulence genes are often found across bacterial ciMGEs and are absent from archaeal ciMGEs. ICEs are by far the main vectors for the accretion of these genes across ciMGEs. I found a total of 9243 virulence genes in 1490 bacterial ciMGEs. Of these 1490 ciMGEs, 1218 correspond to ICEs, 267 IMEs, and only 5 AICEs. As expected, the total number of virulence genes in ICEs also far outnumbers that of IMEs – 8695 and 542, respectively. After correcting the number of virulence genes to the size of the carrying ciMGE, I observed that IMEs in Firmicutes carry more virulence genes per kb than do ICEs (**Supplementary Figure S4A**, p-value = 1.1e-08). I discovered a total of 13 virulence categories across ICEs, dominated by genes related to adherence, exotoxin, immune modulation, and effector delivery system, while out of the 11 virulence categories found on IMEs, the most frequently identified are mostly related to immune modulation, exotoxin, and motility (**Figure 3A** and **Supplementary Table S4**). The 6 virulence genes found on the 5 AICEs from Actinobacteriota are devoted to effector delivery system and as nutritional/metabolic factors.

**Figure 3.**
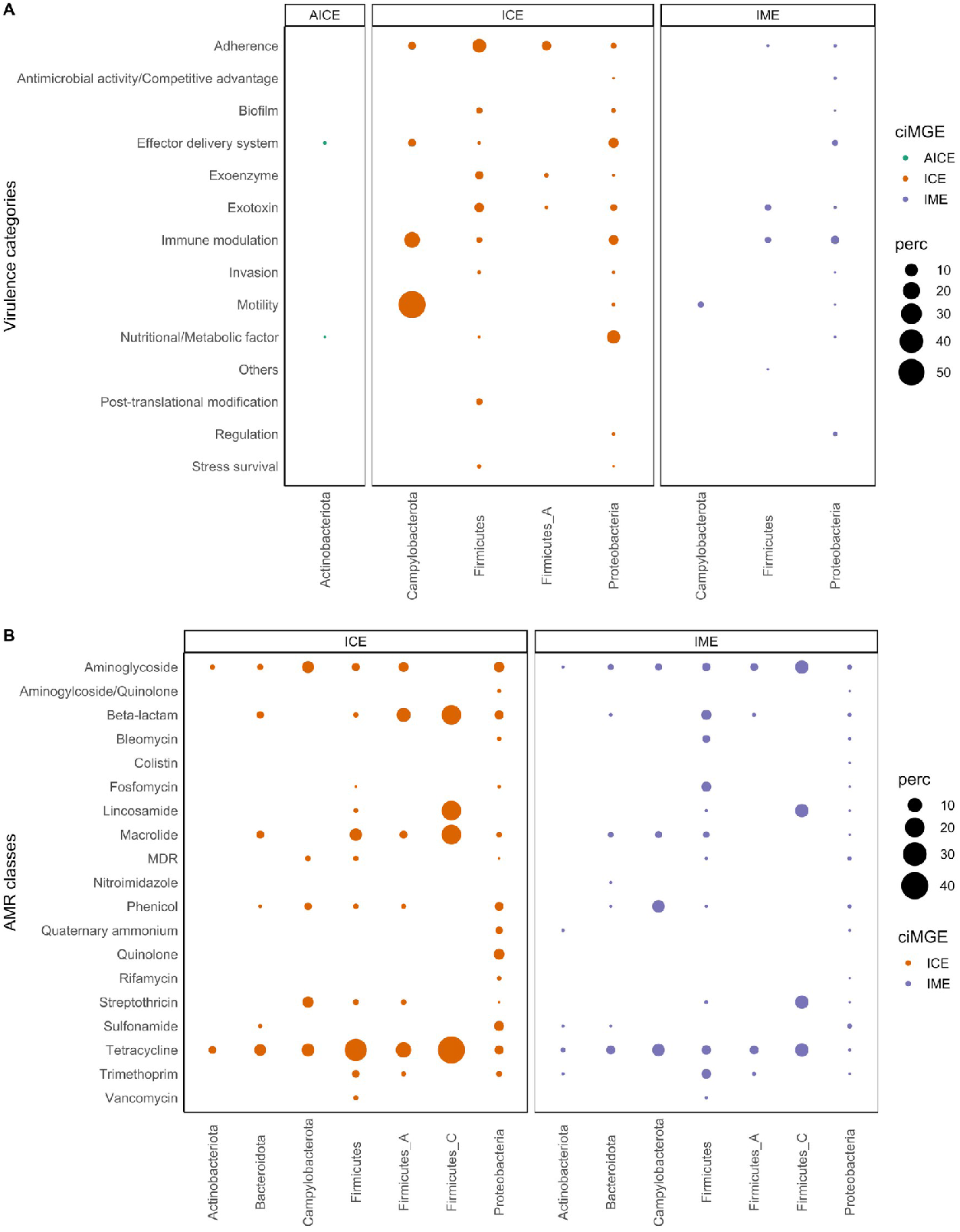
Virulence and AMR genes are more frequently found across ICEs. **A)** Distribution of virulence categories across multiple phyla. Phyla in the x-axis are listed from left to right in alphabetical order. Phyla with ciMGEs carrying genes associated with only one virulence category were excluded. **B)** Distribution of AMR classes across multiple phyla. Phyla in the x-axis are listed from left to right in alphabetical order. Only phyla with more than 5 different AMR classes are shown. The size of the circles is proportional to the percentage of particular virulence categories or AMR classes per ciMGE type per phylum. Circles are coloured according to the ciMGE type.

Known AMR genes are found across bacterial ICEs and IMEs, and absent from archaeal ciMGEs. I discovered a total of 3087 AMR genes in 1241 bacterial ciMGEs. Of these 1241 ciMGEs, 944 correspond to ICEs and 297 to IMEs. This means AMR genes are also more often found in ICEs - 16,1% ICEs carry at least one AMR gene (944/5865), while only 4,7% IMEs carry at least one of these genes (297/6319). The total number of AMR genes was also higher in ICEs than in IMEs - 2394 vs 693, respectively. Curiously, even though the total number of virulence genes in ciMGEs far exceeds that of AMR genes, AMR genes are spread across a wider range of phyla than virulence genes are (12 vs 5, **Supplementary Table S2**). After correcting the number of AMR genes to the size of the carrying ciMGE, I noted that that IMEs in Bacteroidota, Firmicutes, Firmicutes_A, and Proteobacteria carry more AMR genes per kb than do ICEs (**Supplementary Figure S4B**, p-value 1.5e-07, < 2.2e-16, 8.3e-07, and 5.8e-14, respectively). The AMR genes found across ICEs and IMEs encode resistance to a total of 17 and 18 AMR classes, respectively, and most confer resistance to tetracyclines, aminoglycosides, and beta-lactams (**Figure 3B** and **Supplementary Table S5**). Finally, the distribution of defense systems, AMR, and virulence genes was compared between ICEs and IMEs from different phyla. The number of defense system genes is significantly higher than that of AMR and virulence genes across most ICEs and IMEs from multiple phyla (**Supplementary Figure S5**). Whereas the number of AMR genes in ICEs and IMEs is higher than that of virulence genes in Actinobacteriota, Bacteroidota, and Firmicutes_A, virulence genes outnumber AMR genes in both ciMGEs from Proteobacteria (**Supplementary Figure S5**). Taken together, these results underline the role of ICEs as important hotspots for the accumulation of defense systems, virulence, and AMR genes across ciMGEs from multiple phyla.

### Defense systems, AMR, and virulence genes are negatively correlated across ICEs and IMEs

I then explored to what extent the prevalence of defense systems, AMR classes, and virulence categories are correlated across ICEs/IMEs. Since AICEs carry no AMR genes, the analysis was focused exclusively on ICEs and IMEs. Given that the distribution for each class is not normal, the non-parametric Spearman correlation coefficient was used. These three functions are inversely correlated across the ICEs and IMEs identified in this study (**Figure 4**). While these functions are negatively correlated within the same ciMGE, genomes may compensate this correlation by acquiring ciMGEs with the distinct functions. However, I found that defense systems, AMR, and virulence genes are not only correlated within the same ciMGE, but also within the same genome (**Supplementary Figure S6**).

**Figure 4.**
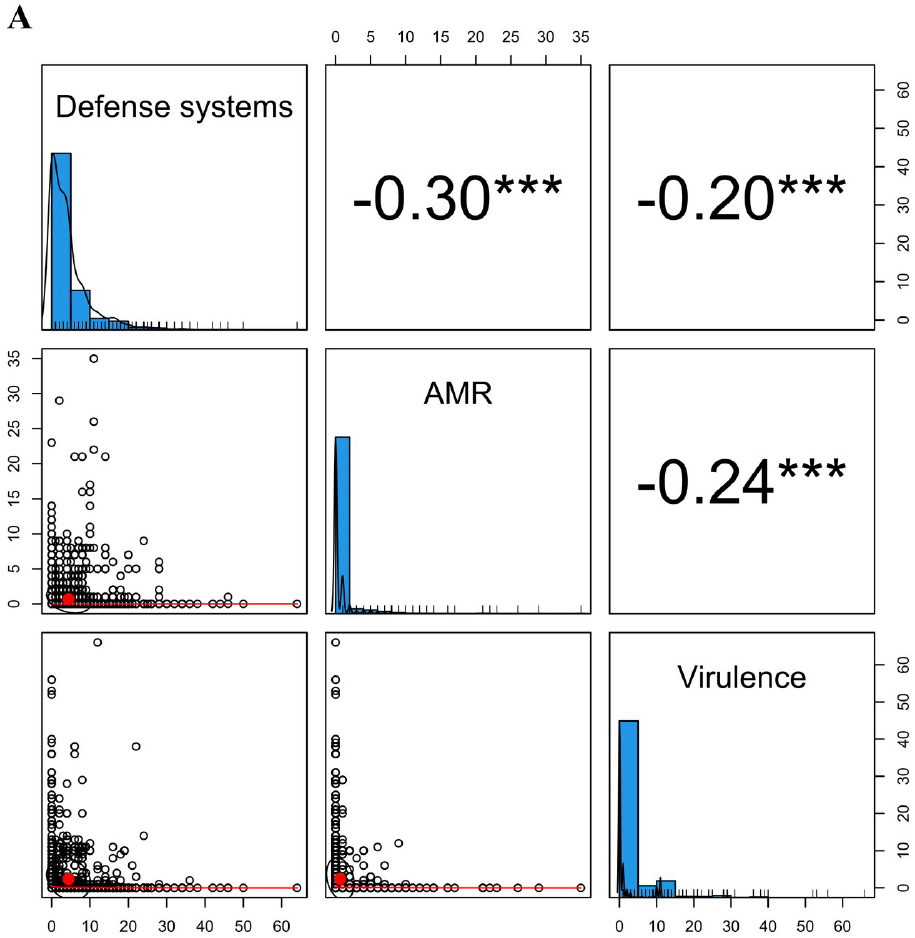
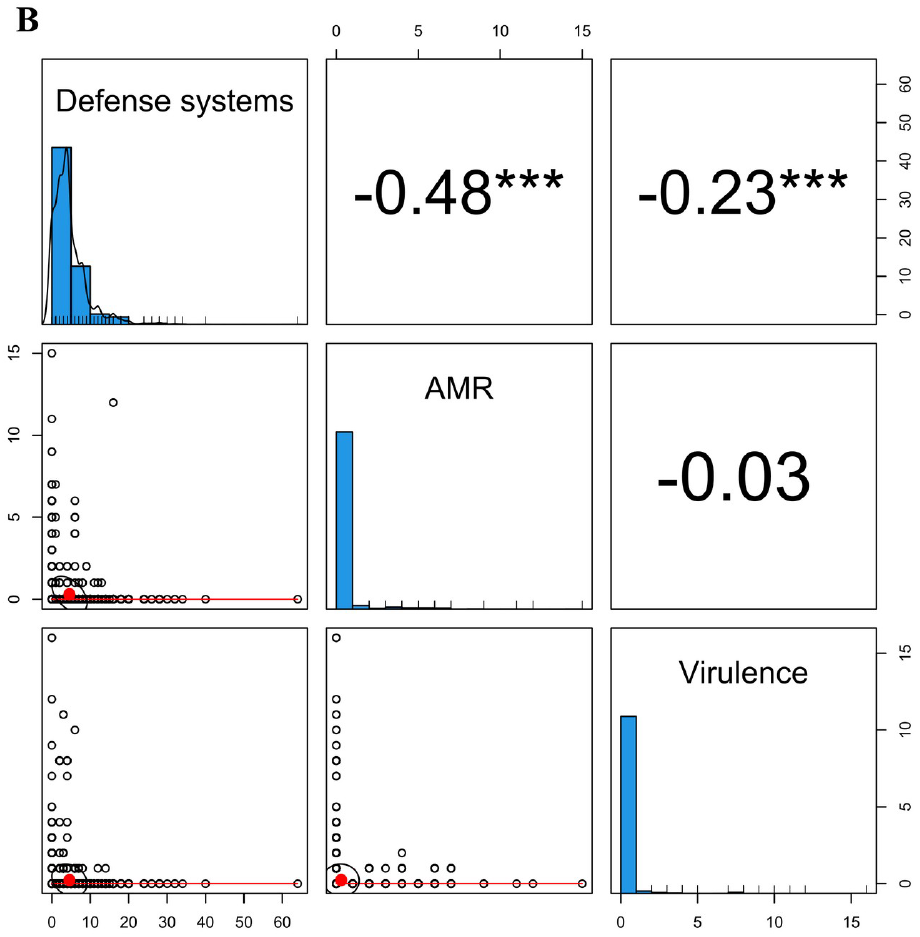
Defense systems, virulence, and AMR genes are negatively correlated. **A)** Correlations across ICEs. **B)** Correlations across IMEs. The scatter plot of matrices shows histograms of distribution of each function on the diagonal, with defense systems in the top left, AMR in the middle, and virulence genes in the bottom right. Scatter plots between pairwise functions are shown below the diagonal, while the Spearman correlation coefficients between functions are shown above the diagonal. Values above 0.05 were considered as non-significant and no asterisks are shown. The following convention was used for symbols indicating statistical significance: * for p <= 0.05, ** for p <= 0.01, *** for p <= 0.001, and **** for p <= 0.0001.

Multiple cases of negative correlations were found in proteobacterial ICEs, such as between virulence genes acting as effector delivery systems and multiple defense systems such as restriction-modification systems and CBASS. Still, positive correlations were observed in specific cases. For example, genes encoding resistance to multiple antibiotics were positive correlated with GAO, BREX, and Abi defense systems across proteobacterial ICEs (**Supplementary Figure S7A**). Positive correlations were also found between virulence genes associated with nutritional/metabolic factors and retrons and DRT defense systems. Curiously, different associations were found across ICEs from Firmicutes (**Supplementary Figure S7B**). For example, Abi and restriction-modification systems were negatively correlated with multiple virulence categories, including exoenzymes and genes involved in adherence functions. Positive correlations were found between tetracyclines and viperins, as well as between virulence genes involved in post-translation modification and Lamassu defense systems. Additionally, genes encoding resistance to multiple antibiotics were positively correlated with Kiwa defense systems.

When looking into proteobacterial IMEs (**Supplementary Figure S7C**), negative correlations were found between multiple defense systems and virulence genes involved in immune modulation. Genes encoding resistance to distinct antibiotic classes (e.g., beta-lactams, aminoglycosides, and sulphonamides) were positively correlated, consistent with the previous observations that these genes tend to be co-localized in the same genetic environment (40). Finally, negative correlations were found between beta-lactams and exotoxins and restriction-modification systems across IMEs from Firmicutes (**Supplementary Figure S7D**). Additionally, Lamassu defense systems were positively correlated with genes encoding resistance to aminoglycosides and virulence genes involved in immune modulation. These results show that while defense systems, AMR, and virulence genes are inversely correlated across ICEs and IMEs, negative and positive correlations between specific genes belonging to these functions can be observed in both ciMGEs.

### ICEs and IMEs form heterogeneous communities composed of multiple phyla

Given the presence of highly similar MGEs in this dataset, the 13274 elements found here were dereplicated into a representative set of 9618 ciMGEs (nucleotide identity threshold of 90% and 80% coverage). Still, focusing on the ciMGEs that clustered above this threshold is an opportunity to explore the presence of highly similar ciMGEs across different taxa (**Supplementary Table S6**). Identical or nearly identical ciMGEs were found in different species, such as similar IMEs in *Prevotella intermedia, Prevotella melaninogenica C, Butyricimonas faecalis, Bacteroides fragilis A* and in *Bacteroides salyersiae* (cluster representative NZ_CP024696_2703150..2736386 in **Supplementary Table S6**). Despite being uncommon, similar IMEs (cluster representative NZ_CP010453_896607..900662) were discovered in three phyla: Firmicutes, Firmicutes_A, and Actinobacteriota.

Each of the 9618 ciMGEs was then reduced to a set of *k*-mers and the Jaccard index (JI) was used as a measure of nucleotide sequence similarity between all ciMGE pairs. Next, I used an alignment-free nucleotide sequence similarity comparison between the representative set of 9618 ciMGE pairs to infer an undirected and weighted network, resulting in 768874 pairwise comparisons. In line with the high diversity frequently observed across MGEs, the majority of ciMGE pairs shared little similarity, with JI values below 0.25 (**Supplementary Figure S8**). As observed for plasmid networks (41), a JI threshold was applied to increase the sparsity of the network and to optimize the performance of the community detection tools Infomap and OSLOM. Using the mean value (0.0374799) of the estimated JI values between the 9618 ciMGEs identified in this study as a threshold, I was able to upweight more similar ciMGEs, resulting in a sparse network of 6424 nodes and 96065 edges. Then, 4935 ciMGEs were grouped into 300 communities (i.e., set of nodes that are more densely connected with one another than expected by chance, **Figures 5A and 5B**), each community including 3 ciMGEs or more. The remaining 1489 ciMGEs formed singletons and pairs and were removed from the analysis. Most communities were phylum-specific, but heterogeneous with respect to the presence of ICEs and IMEs. For example, AICEs from Actinobacteria were mostly clustered in homogeneous communities, while ICEs and IMEs typically formed heterogeneous communities dominated by either Proteobacteria, Firmicutes, Bacteroidota or Campylobacterota. The absence of pairwise distance similarities with intermediate JI (**Supplementary Figure S8**) helps to explain this clustering in discrete communities, instead of a continuous genetic structure. Remarkably, these ciMGEs communities were not homogeneous to a given relaxase family. The relaxase MOB_T_ family was the most frequently identified across this set of 4935 ciMGEs, followed by MOB_P1_ and MOB_H_ (**Supplementary Table S7**). Altogether, these results underline the broad host range of ICEs and IMEs and their ability to challenge interphylum barriers and to form heterogeneous communities with elements from multiple phyla.

**Figure 5.**
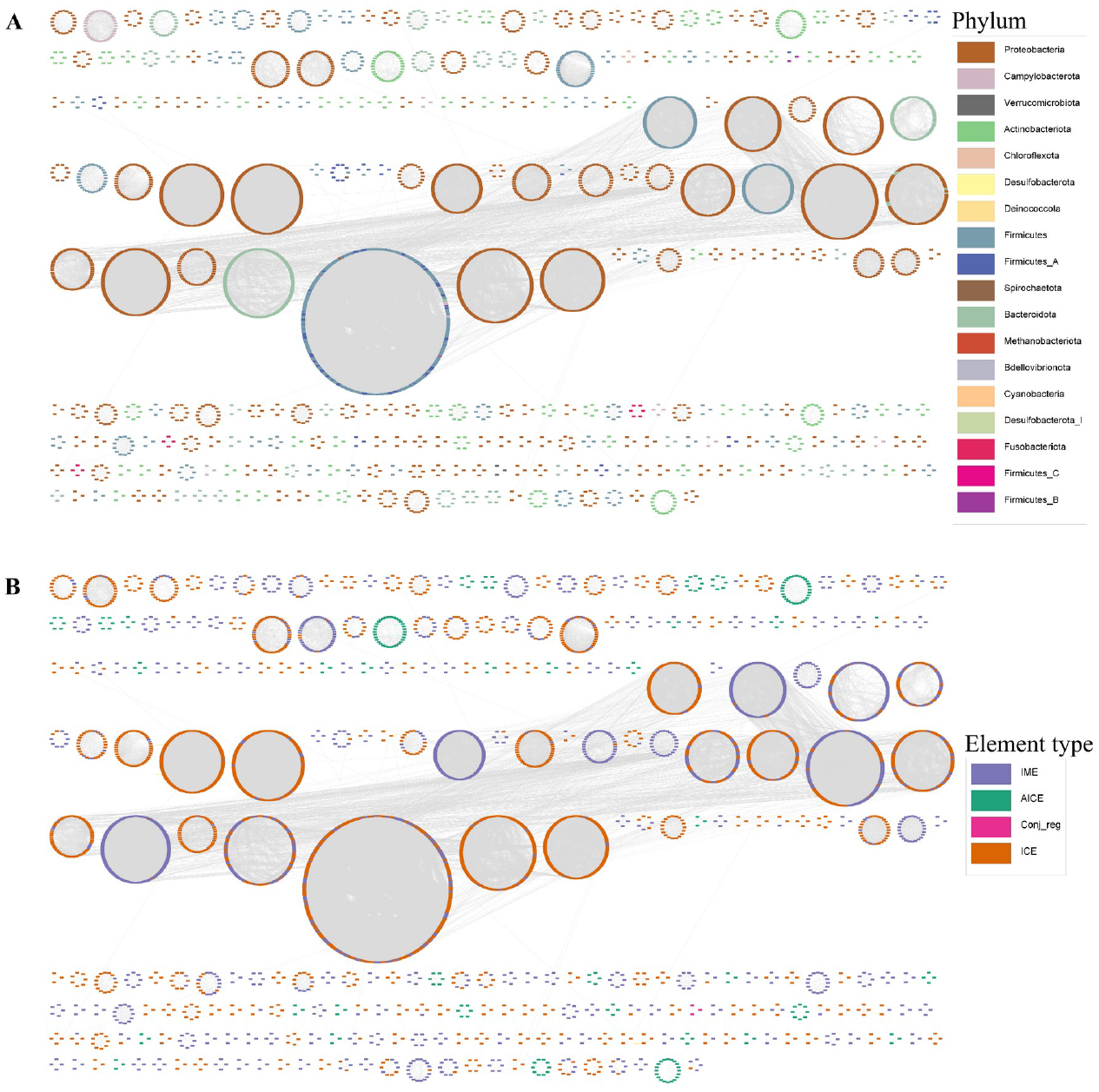
Weighted network of ciMGE communities detected with Infomap and Oslom. Nodes coloured by **A)** Phylum; and **B)** ciMGE type. Each ciMGE is represented by a node, connected by edges according to the Jaccard index value between all ciMGE pairs. Singletons and pairs were excluded from the network.

### The functional landscape of ICEs is populated by uncharacterized proteins

The proteome of the dereplicated dataset of ciMGEs from both Bacteria and Archaea (total number of proteins = 629300) was then scanned for cluster or orthologous groups (COGs) categories. Given that ICEs are usually larger than IMEs (**Figure 1D**), the absolute counts of COG categories found on each ICE and IME (**Supplementary Table S8**) was corrected to the size of the corresponding ICE and IME, respectively. For the majority of COG categories, the normalized number of IME proteins with an assigned function was significantly higher than that of ICE proteins (p<2.2e-16, **Figure 6**). This difference was consistently observed across the phyla with a higher incidence of both ICEs and IMEs: Proteobacteria, Firmicutes, Actinobacteriota, Bacteroidota, Campylobacterota, and Firmicutes_A (**Supplementary Figure S9**). Interestingly, the only COG category that was significantly more assigned on ICE proteins was the category S, which refers to unknown functions (p<2.2e-16, **Figure 6**).

**Figure 6.**
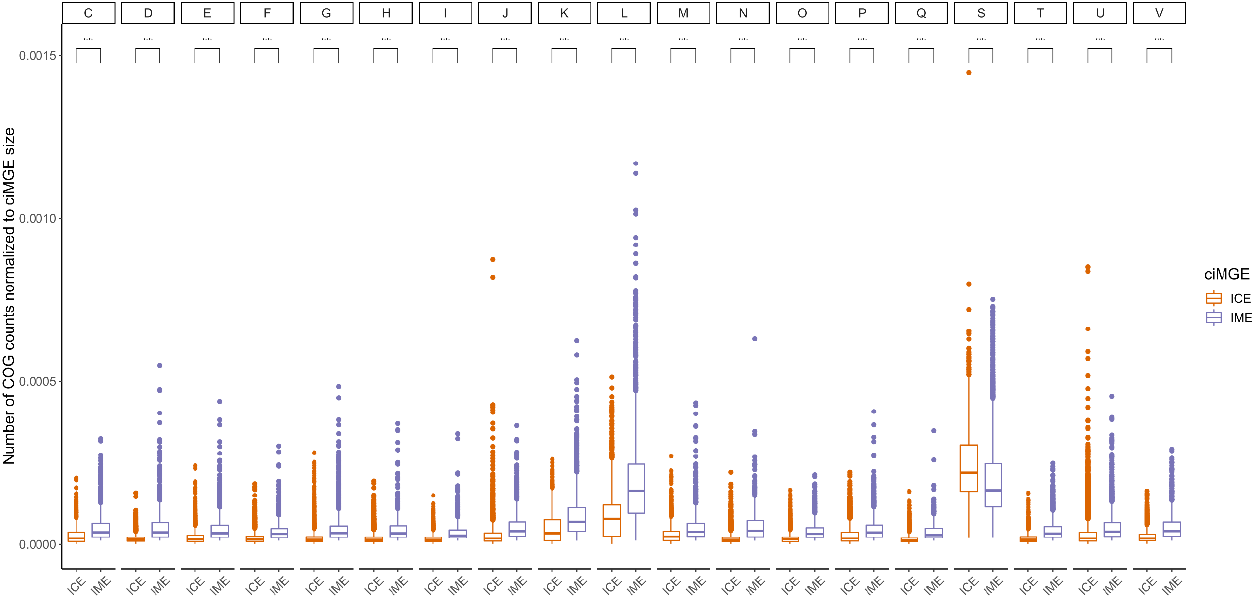
The normalized number of unknown functions is significantly higher across ICEs. Boxplots showing the number of COG categories found in each ICE or IME from the six major phyla found in this study and normalized to the size of the ICEs and IMEs. Comparisons were performed using the Wilcoxon test, and the p-values adjusted with the Holm–Bonferroni method. The following convention was used for symbols indicating statistical significance: * for p <= 0.05, ** for p <= 0.01, *** for p <= 0.001, and **** for p <= 0.0001. Boxplots are coloured according to the ciMGE tpye. COG categories: C - Energy production and conversion; D - Cell cycle control, cell division, chromosome partitioning; E - Amino acid transport and metabolism; F - Nucleotide transport and metabolism; G - Carbohydrate transport and metabolism; H - Coenzyme transport and metabolism; I - Lipid transport and metabolism; J - Translation, ribosomal structure and biogenesis; K – Transcription; L - Replication, recombination and repair; M - Cell wall/membrane/envelope biogenesis; N - Cell motility; O - Posttranslational modification, protein turnover, chaperones; P - Inorganic ion transport and metabolism; Q - Secondary metabolites biosynthesis, transport and catabolism; S - Function unknown; T - Signal transduction mechanisms; U - Intracellular trafficking, secretion, and vesicular transport; V - Defense mechanisms. COG categories A (RNA processing and modification), B (Chromatin structure and dynamics), W (Extracellular structures), and Z (Cytoskeleton) were excluded due to low COG counts.

It was also found that a higher relative proportion of IME proteins have known functions across the most common bacterial phyla identified in this study (**Figures 6** and **S7**). On the other hand, ICEs have more uncharacterized proteins, which suggests the existence of a pool of unknown genes with functions yet to be discovered. Curiously, no hits for COG category X (mobilome: prophages, transposons) were found in this study. Since ICEs are also known as conjugative transposons and share genetic features with prophages (such as the presence of phage integrases), it would be expected to find protein hits for this COG category. This can be explained by the fact that COGs assigned to both the L and X categories in the NCBI’s Database of COGs (for example, COG0582 – integrase/recombinase) were solely assigned to the L category using eggNOG-mapper. In fact, COG0582 was the most commonly observed COG across category L, which helps to explain the high values for the normalization counts of this category across this dataset. A similar trend was observed for insertion sequences and transposases, which are common across ciMGEs and were here assigned to category L.

## Discussion

In this study, I characterize a collection of more than 13000 ciMGEs, including more than 6000 IMEs, nearly 6000 ICEs, and more than 1000 AICEs. AICEs are frequently seen across Actinobacteriota, however in this study one element was detected in Proteobacteria and four elements in Firmicutes (**Supplementary Tables S2 and S9**). Comparing with the number of ciMGEs with nucleotide sequences available at ICEberg 2.0 (https://bioinfo-mml.sjtu.edu.cn/ICEberg2/download.html), this work represents a massive increase in the number of publicly available IMEs (6331 vs 111), ICEs (5869 vs 718), and AICEs (1061 vs 50). These large discrepancies are somehow anticipated, since the number of ciMGEs available at ICEberg was last updated in 2018, and the number of bacterial and archaeal genomes have been skyrocketing ever since (42). On top of that, ciMGEs characterized in this study were corrected for taxonomy of the host genome, based on the rank-normalized classification module from domain to species available at the Genome Database Taxonomy (43). The distribution of a ciMGE’s sequence length depends on whether it is an IME, ICE, or AICE, while its GC content is closely tied to that of its host. The left-skewed distribution observed for AICE’s GC content can be explained by the high GC content of the bacterial host. In fact, the vast majority of AICEs were identified in Actinobacteriota, which typically have high GC content (23). On the other hand, the multimodal distribution of ICEs/IMEs’ GC content can be explained by the wide distribution of these elements across multiple phyla with variable GC content.

Recently, it was shown that mobility genes (such as transposon and prophage components) and defense systems are non-randomly clustered in genomic islands (2), but the type of ciMGEs involved in this process was not assessed. Here, I show that a wide repertoire of defense systems is accumulated across ICEs and IMEs from multiple phyla. Crucially, I found that defense systems, AMR genes, and virulence genes are inversely correlated across bacterial ICEs and IMEs, suggesting that carrying multiple cargo genes with distinct functions in the same ciMGE is detrimental to bacterial fitness. An increased MGE size is one of the primary mechanisms by which numerous cargo genes can be harmful. Larger MGEs may be harder for bacteria to move amongst one another and may impose a higher metabolic burden on the host cell (44), leading to lower fitness of the host cell, especially under conditions of stress or resource limitation. Overall, the presence of multiple cargo genes in MGEs can have both positive and negative effects on the fitness of the host bacteria, with the balance between these effects depending on the specific genetic and environmental context While defense systems and AMR genes were identified across ciMGEs from multiple phyla, virulence genes were limited to five phyla (**Figure 3A**). This can in part be explained by the inclusion of only 32 genera of pathogens with medical importance in VFDB with full information available (32), meaning that virulence genes from multiple phyla were most likely missed from this analysis.

The network-based approach used in this work revealed that most ciMGEs form clusters of high nucleotide identity that are homogeneous to the host phylum, which is in agreement with the phylogenetic and biological barriers that shape HGT events (45). Still, distantly related phylum interactions were observed across multiple ciMGEs, for example between Campylobacterota, Actinobacteriota, and Firmicutes (**Figure 5A**), in line with the broad host range attributed to these elements (46). Interestingly, the dereplication approach used to build the ciMGE network (using a 90% nucleotide identity threshold) removed no elements from Archaea, uncovering that ciMGEs in this domain do not share high nucleotide identity. Indeed, only two elements (of the 16 ciMGEs found in archaeal genomes from this dataset) are plotted in the network and form a small cluster exclusively containing these ciMGEs (**Figure 5A**), meaning the JIs including most archaeal ciMGEs fall below the mean threshold for all pairwise comparisons between Bacteria and Archaea, and exposing how distantly related these elements are.

There are several bioinformatic tools available to search for extrachromosomal elements such as plasmids, however there are currently few alternatives for ciMGEs as ICEs and IMEs. Recently, ICEscreen was developed (47), with the purpose of detecting these elements across Firmicutes. I decided to use ICEfinder (25), since it is not restricted to a particular phylum. Yet, it should be noted that this tool may not provide the precise boundaries for ciMGEs, as it relies in the 3’ termini of the tRNA/tmRNA genes as insertion sites, even though other integration hotspots can exist, such as into the 3’ or 5’ ends of protein-coding genes. Even though this tool was designed to scan ciMGEs across bacterial genomes, it was able to identify 16 elements in archaeal genomes. Still, it is possible some elements may have been overlooked, since integrases, relaxases, and other signature proteins from bacterial ciMGEs may be too distantly related to those from Archaea. Conceptually, prophages would fit the idea of a chromosomally integrated MGE but were not considered in this study because multiple DNA sequences of these elements are already available in public databases. Despite the fact that the number of publicly available CIMEs deposited in ICEberg is lower than that of ICEs, IMEs, and AICEs (31 vs 718, 111, and 50, respectively), still it is somewhat surprising that no CIMEs were discovered in this study. One possible explanation for this is that these elements frequently form composite structures with ICEs and are often integrated in the same site (48). This tandem accretion may result in incomplete predictions. The recently developed ICEscreen (47) exhibited a good performance in disentangling these complex structures in Firmicutes.

Whereas the integration of plasmids into the chromosome is not the trend in most plasmids, it is common in some cases (ref). Using PlasmidFinder, a total of 372 hits were found across the 13274 ciMGEs identified in this study, mostly in Firmicutes (**Supplementary Table S10**). However, it is important to highlight that although the integration/excision module is traditionally associated with ciMGEs and the replication module with plasmids, ciMGEs may carry genes coding for replicases, while some plasmids may also carry genes encoding integrases (46). As so, these hits may indeed correspond to bona fide ciMGEs instead of plasmid artifacts. This study only focused on genomes that were sequenced at the complete level, meaning the prevalence of ciMGEs across genomes sequenced at the scaffold and contig level was not assessed. Even though genomes sequenced at this level exceed the number of complete genomes by orders of magnitude (42), focusing on the latter was crucial to accurately delineate ICEs and IMEs across ‘intact’ chromosomes. Following the same rationale, metagenome-assembled genomes were also not included in this study. Still, studying the distribution of ICEs and IMEs in relevant reservoirs such as the human microbiome is crucial to better understand the eco-evolutionary dynamics that shape the acquisition of defense systems, AMR, and virulence genes in complex communities (49).

To conclude, this work represents a massive increase in the number of ciMGEs currently available in public databases (from <1000 to >13000). I found that IMEs outstrip ICEs, and both are prevalent across different phyla. ICEs represent the most important ciMGEs for the accumulation of defense systems, virulence genes, and antimicrobial resistance genes. Furthermore, I discovered that these genes are inversely correlated not only within ICEs and IMEs, but also in ICEs and IMEs within the same genome. This suggests that defense systems, AMR, and virulence genes do not tend to coexist in ciMGEs carrying distinct functions within the same genome. Multiple representatives of these two elements share high genetic similarity and challenge phylogenetic barriers. Overall, I discovered that the functional landscape of ICEs is populated by proteins with unknown functions. Finally, this study offers a thorough catalog of ciMGEs from 34 different phyla in the bacterial and archaeal domains, including their nucleotide sequence and associated metadata.

## Data availability

Analyses were made with a combination of shell and RStudio v2022.07.1 scripting. Code used to reproduce major analysis, figures, and supplementary files (including the eggnog output with the functional annotation for the ciMGEs proteins) are available at the Gitlab repository https://gitlab.gwdg.de/botelho/ices_imes. The nucleotide sequences and associated metadata of the ciMGEs identified in this study are available at the Figshare project https://doi.org/10.6084/m9.figshare.21583413.v1.

## Acknowledgments

I would like to thank Jaime Iranzo and Adrian Cazares for relevant discussions while preparing this manuscript.

