## Supplementary figures S1-S9 for "Defense systems are pervasive across chromosomally integrated mobile genetic elements and are inversely correlated to virulence and antimicrobial resistance"

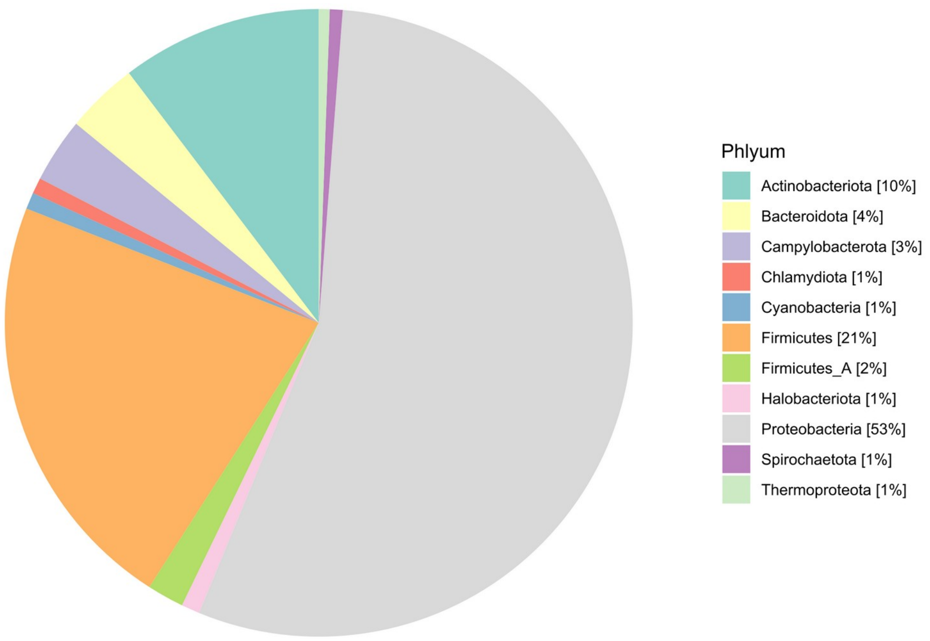


**Figure S1.** Pie chart showing the proportion of genomes used in this study that belong to a particular phylum, according to the GTDB taxonomy. The relative values are shown in percentages next to each phylum in the colour legend. Only phyla with relative values ≥1 are shown in the figure.

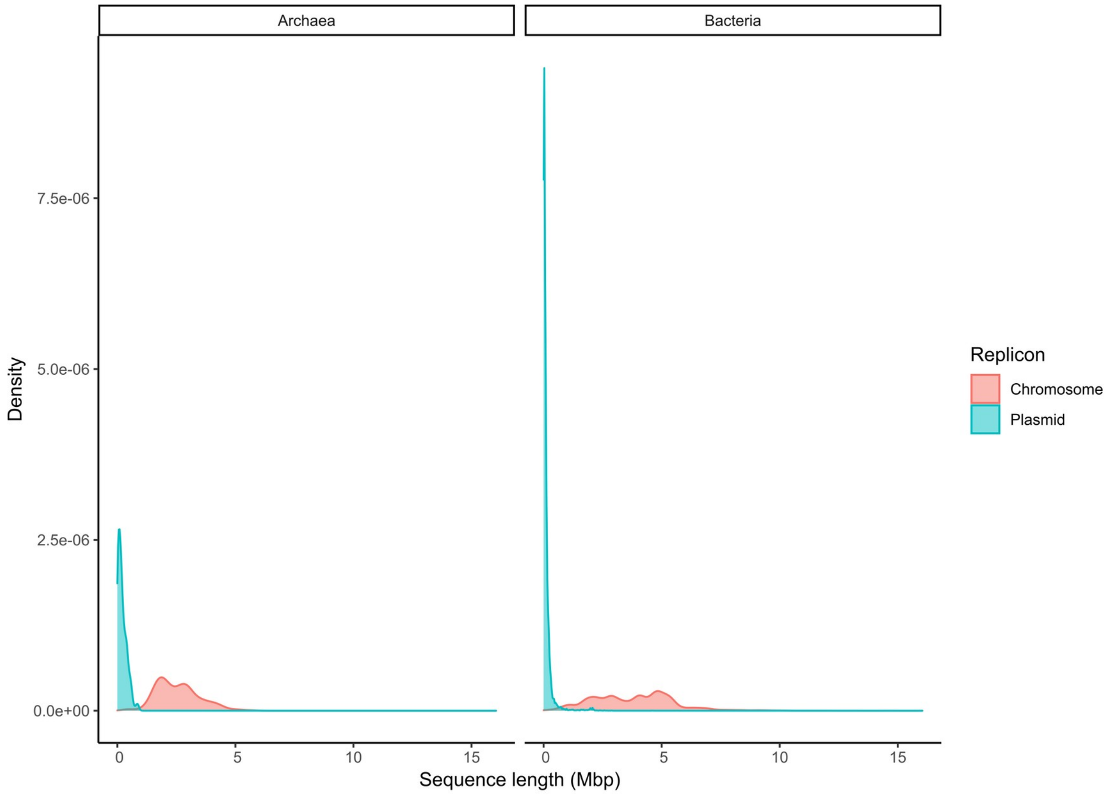


**Figure S2.** Density plot showing the distribution of sequence length (Mbp) for all chromosomal and plasmid replicons used in this study. The left panel shows the distribution of sequence length for archaeal replicons, while the right panel shows the distribution for bacterial replicons.

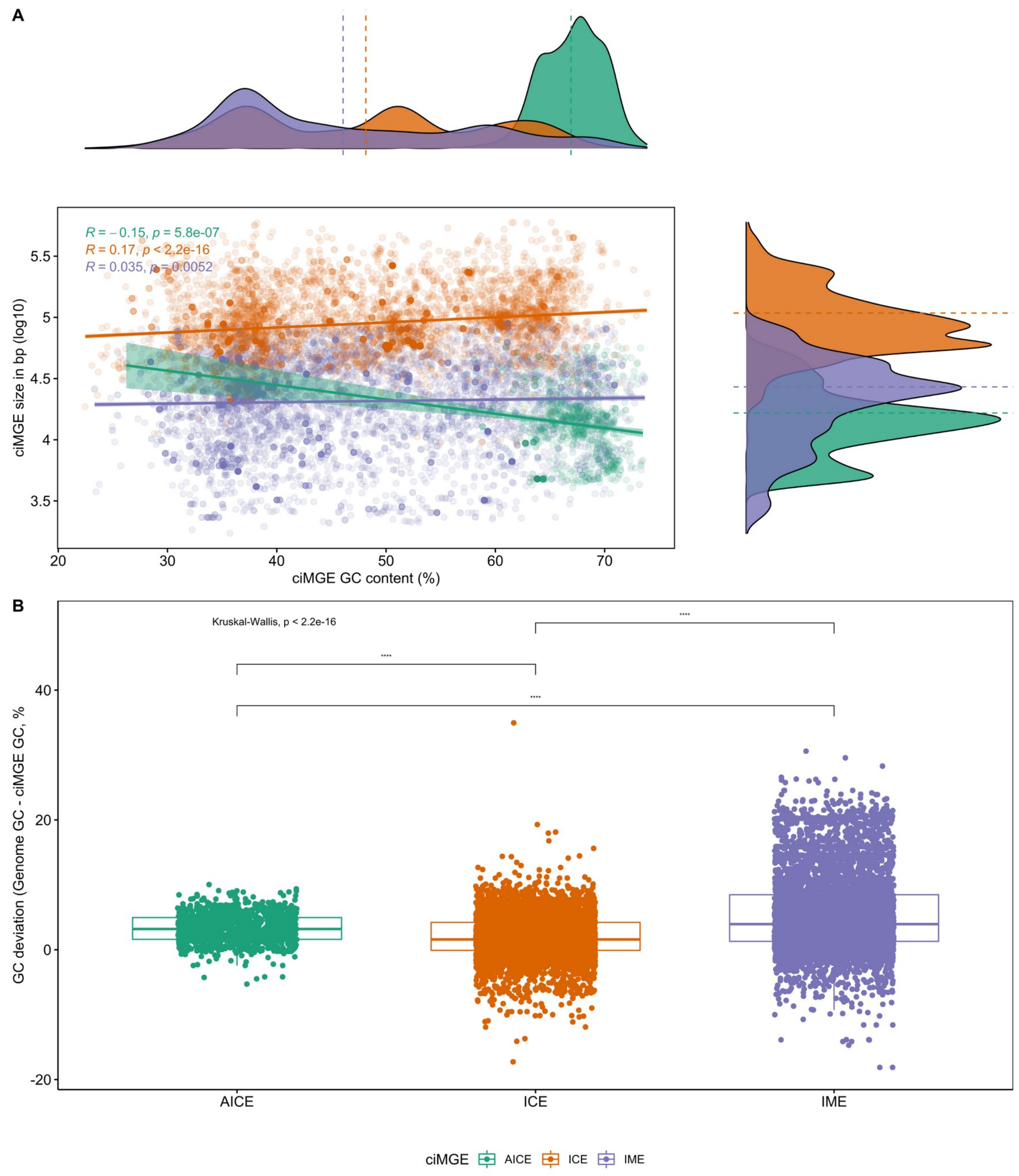


**Figure S3. A)** Scatter plot with marginal density plots comparing the ciMGE size in bp and log10 scaled and the ciMGE GC content (%). Mean values for each ciMGE type are represented by dashed lines in each density plot. **B)** Boxplots showing the GC content variation (i.e., the difference between the genome GC content and that of the ciMGE) across AICEs, IMEs, and ICEs. The following convention was used for symbols indicating statistical significance: * for p <= 0.05, ** for p <= 0.01, *** for p <= 0.001, and **** for p <= 0.0001. Boxplots, density curves, regression lines, dashed lines, and correlation coefficients are coloured according to the ciMGE type.

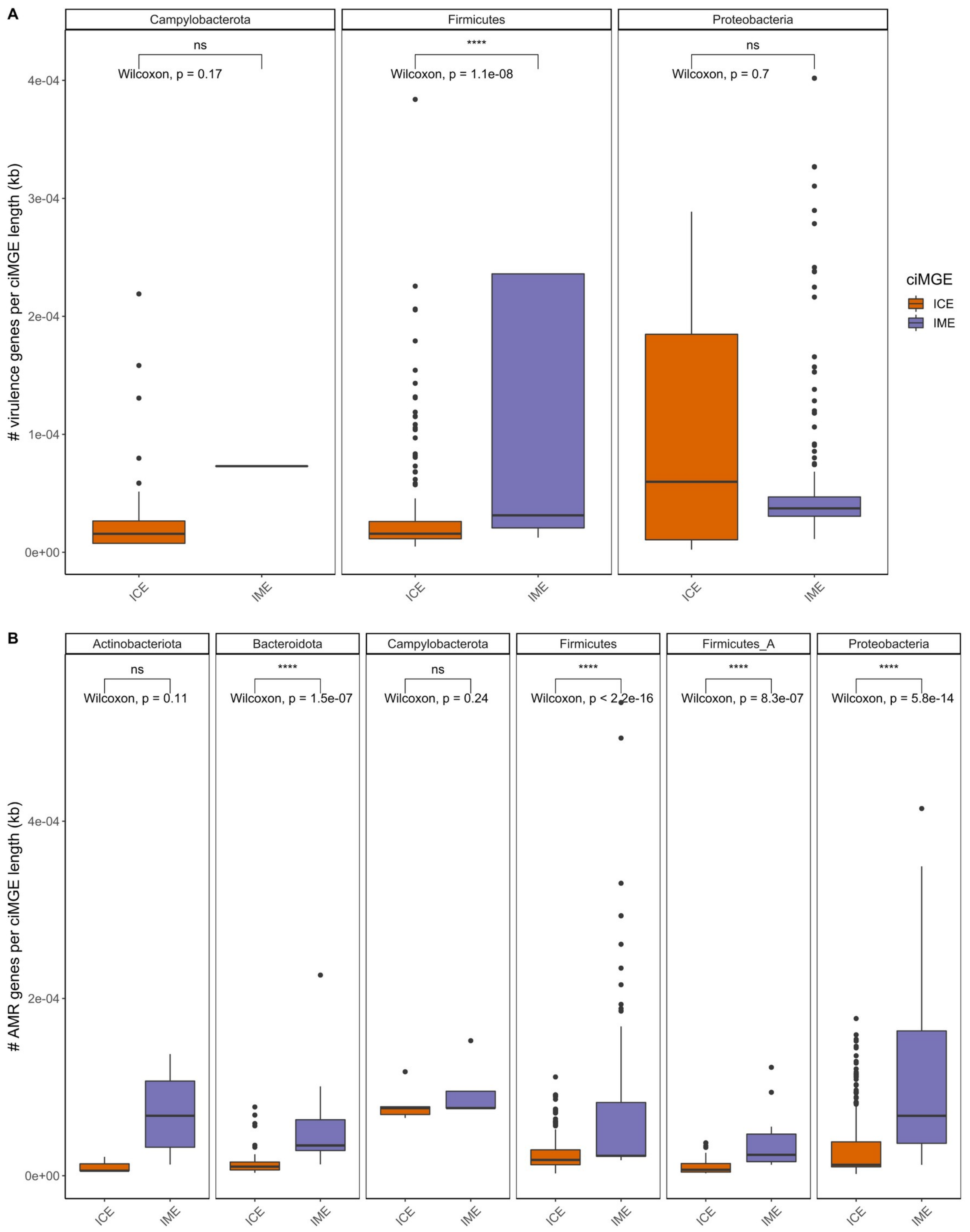


**Figure S4. A)** Boxplots showing the variation between the number of virulence genes normalized by the size (kb) of ICEs and IMEs. Only phyla with more than 2 ciMGEs carrying virulence genes are shown. **B)** Boxplots showing the variation between the number of AMR genes normalized by the size (kb) of ICEs and IMEs. Only phyla with more than 5 ciMGEs carrying AMR genes are shown. Only statistically significant comparisons are shown above the boxplots. The following convention was used for symbols indicating statistical significance: *

for p <= 0.05, ** for p <= 0.01, *** for p <= 0.001, and **** for p <= 0.0001. Boxplots are coloured according to the ciMGE type.

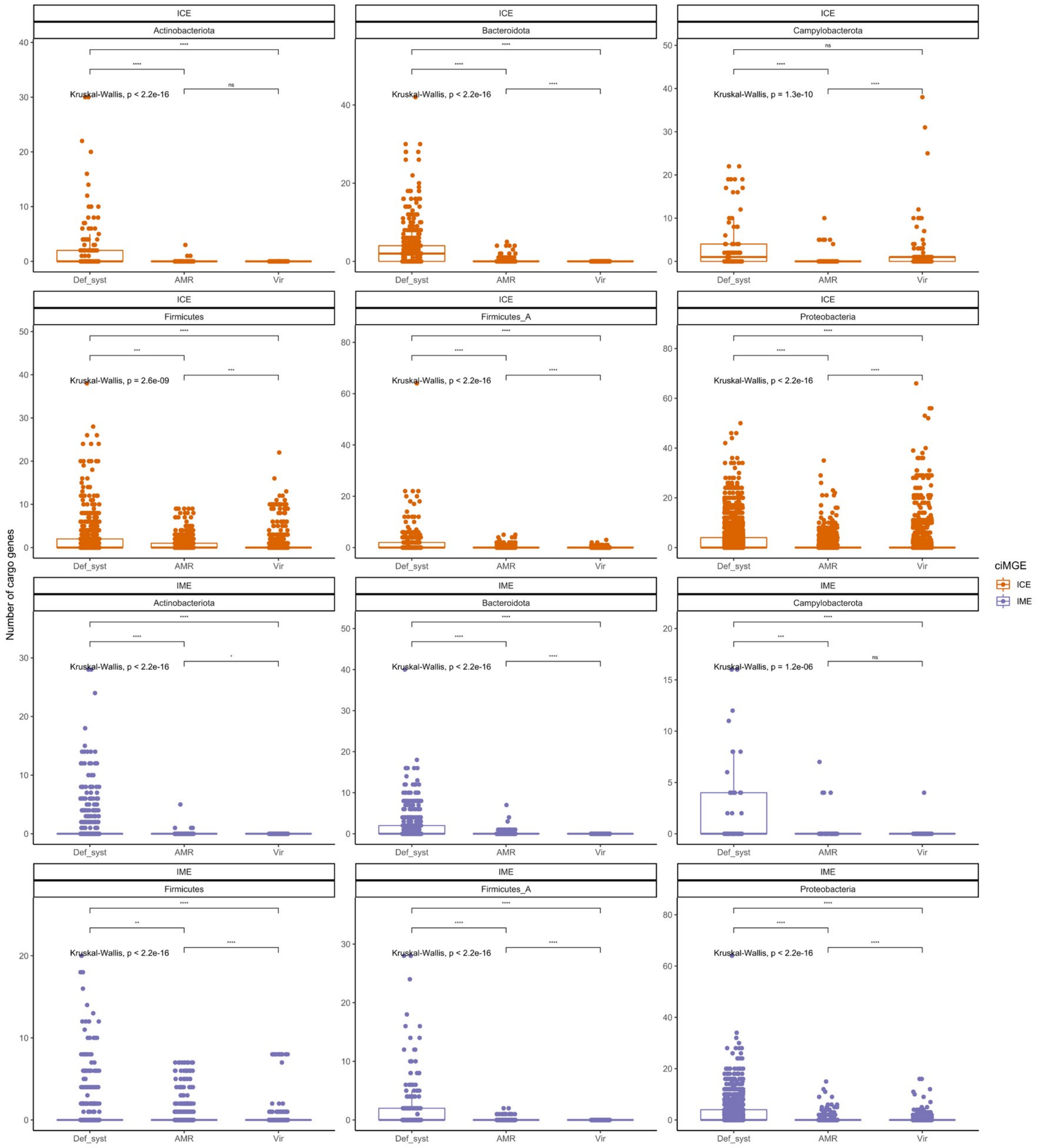


**Figure S5.** Boxplots showing the variation in the number of cargo genes (i.e., defense systems, AMR, and virulence genes) across ICEs and IMEs from multiple phyla. Only phyla with at least 40 IMEs and 40 ICEs are shown. Values above 0.05 were considered as non-significant (ns). The following convention was used for symbols indicating statistical significance: * for p <= 0.05, ** for p <= 0.01, *** for p <= 0.001, and **** for p <= 0.0001. Boxplots are coloured according to the ciMGE type.

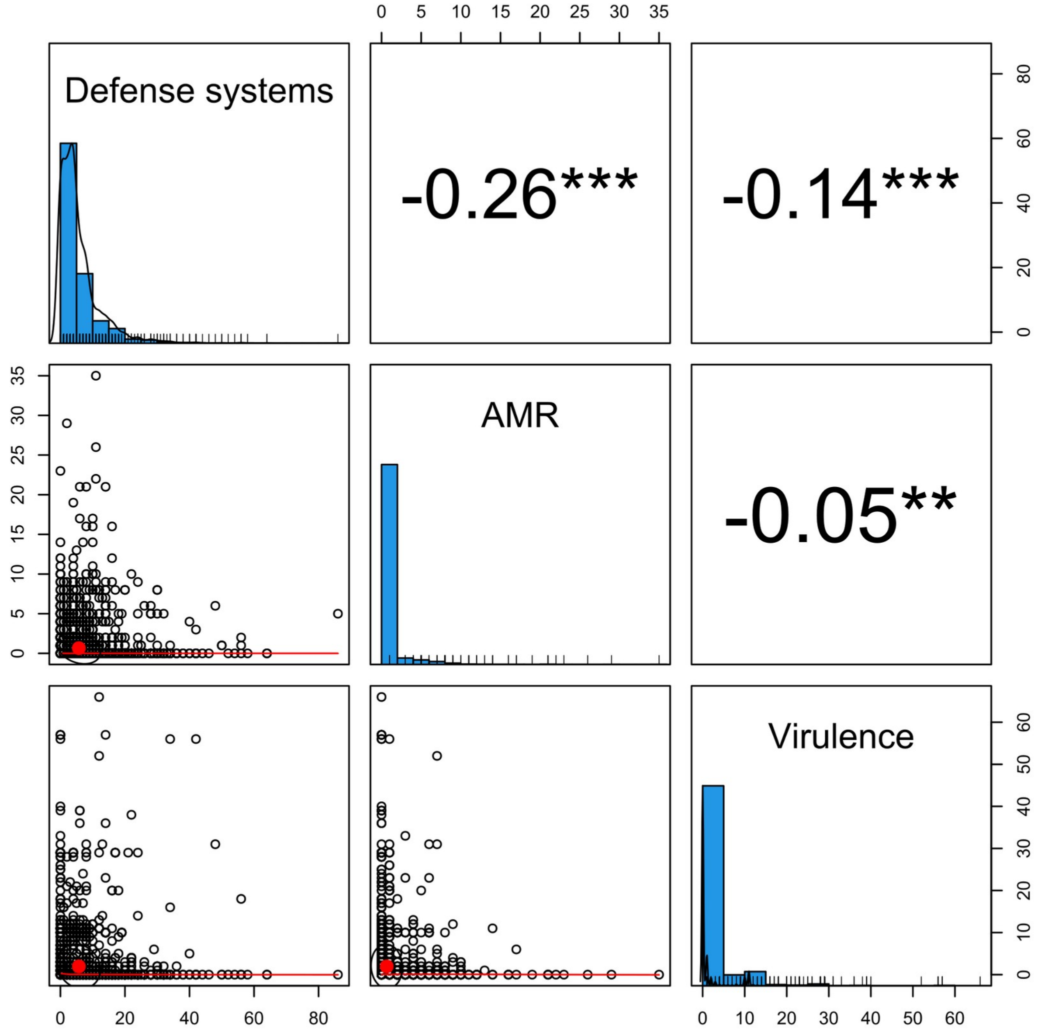


**Figure S6.** Scatter plot of matrices between defense systems, AMR, and virulence genes found within ciMGEs in the same genome. Histograms of distribution of each function are shown on the diagonal, with defense systems in the top left, AMR in the middle, and virulence genes in the bottom right. Scatter plots between pairwise functions are shown below the diagonal, while the Spearman correlation coefficients between functions are shown above the diagonal. Values above 0.05 were considered as non-significant and no asterisks are shown. The following convention was used for symbols indicating statistical significance: * for p <= 0.05, ** for p <= 0.01, *** for p <= 0.001, and **** for p <= 0.0001.

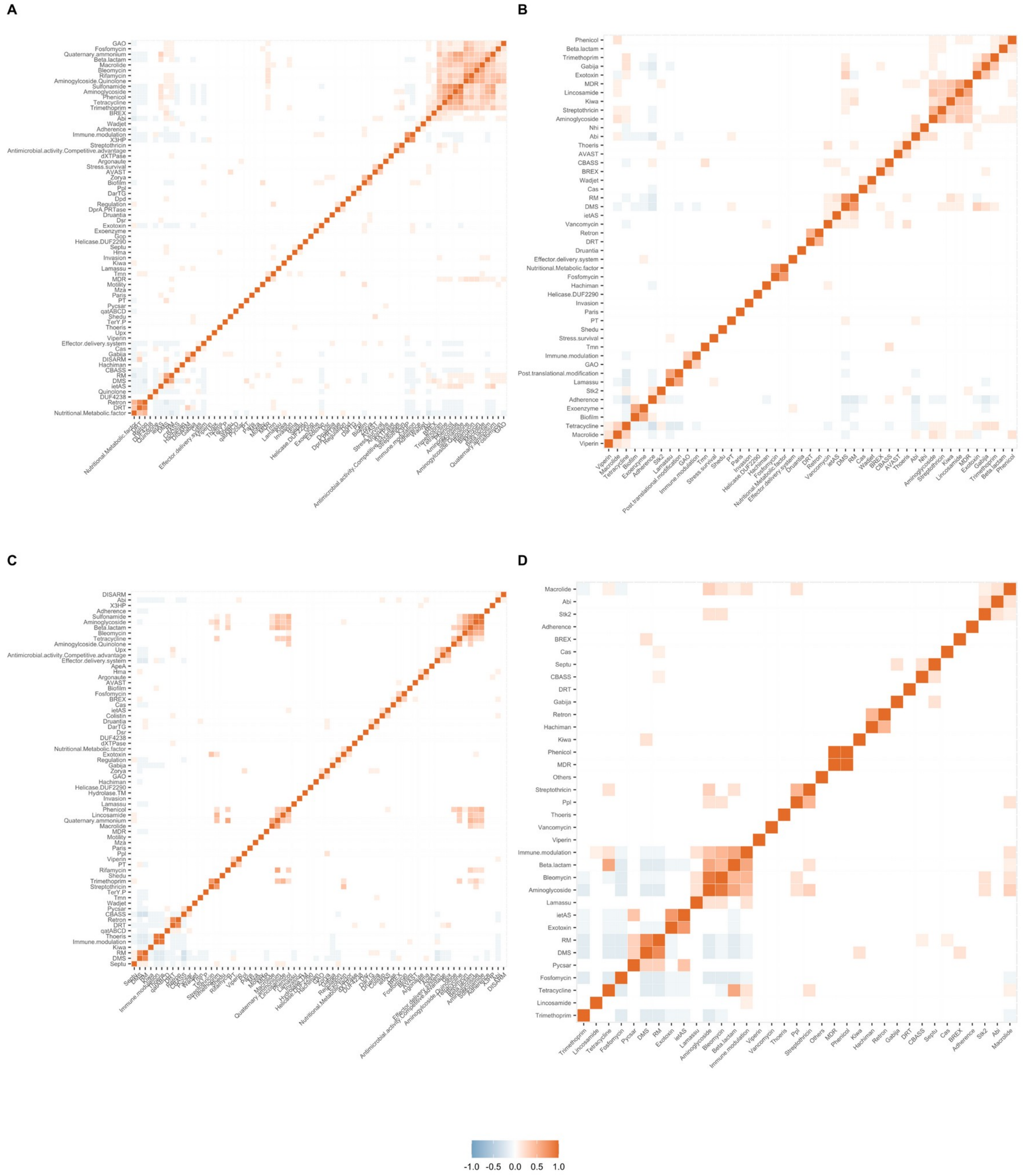


**Figure S7.** Correlation plots between AMR classes, virulence categories, and defence systems across ICEs from Proteobacteria (**A**) and Firmicutes (**B**), and IMEs from Proteobacteria (**C**) and Firmicutes (**D**). The distribution of genes across ICEs and IMEs was converted into a presence/absence matrix. Correlation matrices were ordered using the hierarchical clustering function. Positive correlations are shown in different shades of red, while negative correlations are shown in different shades of blue.

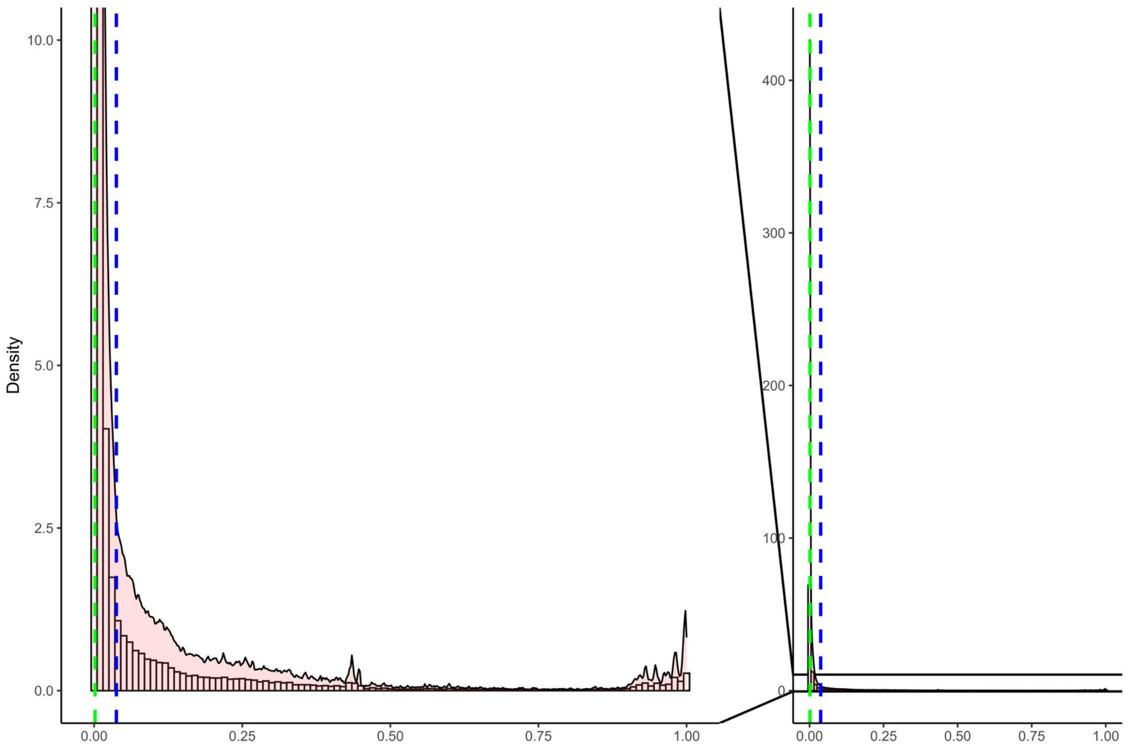


**Figure S8.** Density plot showing the distribution of Jaccard index (JI) values for all versus all pairwise comparisons between the ciMGEs identified in this study. The mean distribution for the JI is shown in blue, while the median is shown in green. Density values from 0-10 were zoomed in and shown in the left. Lines connecting the left and right plots are meant to indicate that the left plot zooms in on the right plot.

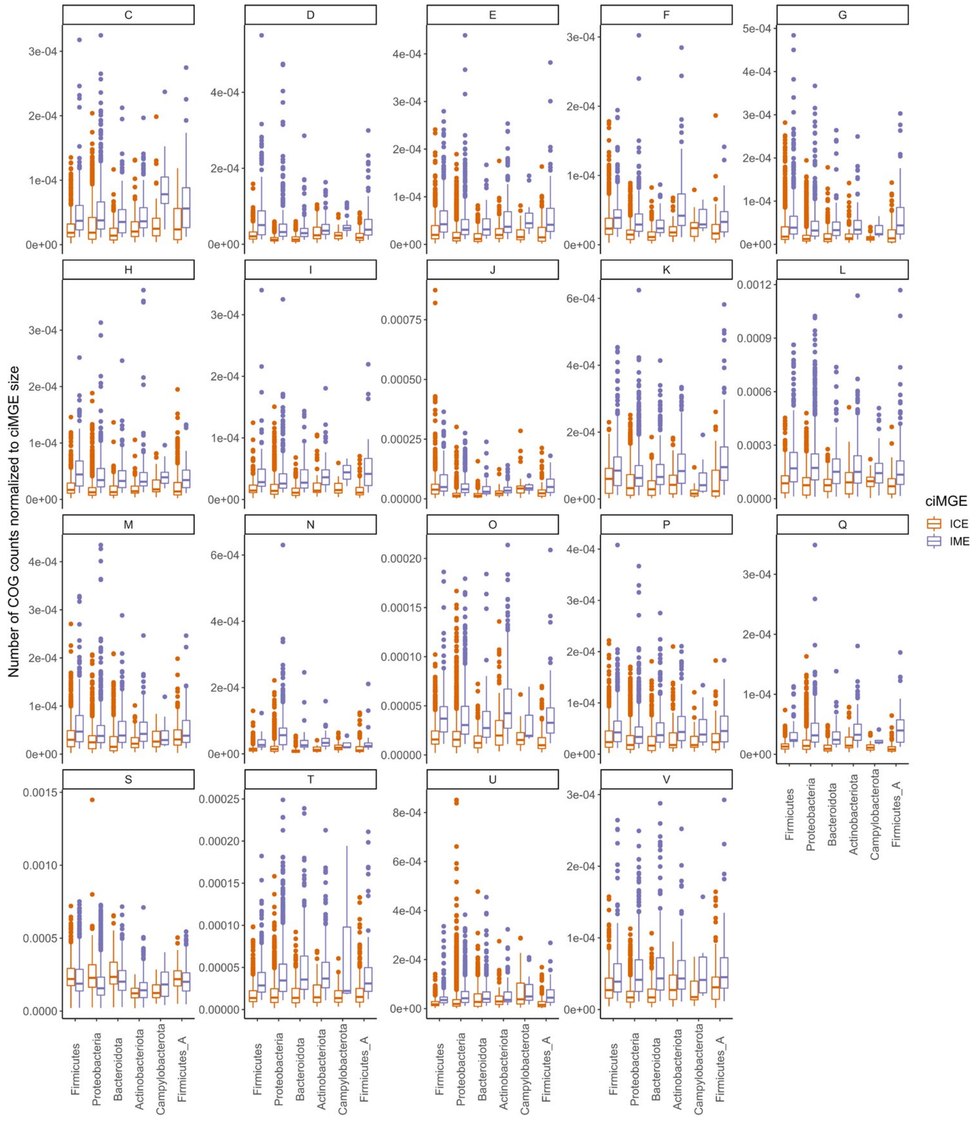


**Figure S9.** Boxplots showing the variation in the COG counts normalized to ICEs and IMEs' size across multiple phyla. Only the six major phyla found in this study are shown. Boxplots are coloured according to the ciMGE type.COG categories: C - Energy production and conversion; D - Cell cycle control, cell division, chromosome partitioning; E - Amino acid transport and metabolism; F - Nucleotide transport and metabolism; G - Carbohydrate transport and metabolism; H - Coenzyme transport and metabolism; I - Lipid transport and metabolism; J - Translation, ribosomal structure and biogenesis; K – Transcription; L - Replication, recombination and repair; M - Cell wall/membrane/envelope biogenesis; N - Cell motility; O - Posttranslational modification, protein turnover, chaperones; P - Inorganic ion transport and metabolism; Q - Secondary metabolites biosynthesis, transport and catabolism; S - Function unknown; T - Signal transduction mechanisms; U - Intracellular trafficking, secretion, and vesicular transport; V - Defense mechanisms. COG categories A (RNA processing and modification), B (Chromatin structure and dynamics), W (Extracellular structures), and Z (Cytoskeleton) were excluded due to low COG counts.
